# Word detection in individual subjects is difficult to probe with fast periodic visual stimulation

**DOI:** 10.1101/2020.10.27.355032

**Authors:** Lydia Barnes, Selene Petit, Nicholas Badcock, Christopher Whyte, Alexandra Woolgar

## Abstract

Measuring cognition in single subjects presents unique challenges. Yet individually sensitive measurements offer extraordinary opportunities, from informing theoretical models to enabling truly individualised clinical assessment. Here, we test the robustness of fast, periodic, visual stimulation (FPVS), an emerging method proposed to elicit detectable responses to written words in the electroencephalogram (EEG) of individual subjects. The method is non-invasive, passive, and requires only a few minutes of testing, making it a potentially powerful tool to test comprehension in those who do not speak or who struggle with long testing procedures. In an initial study, Lochy et al. (2015) used FPVS to detect word processing in 8 out of 10 fluent French readers. Here, we attempted to replicate their study in a new sample of ten fluent English readers. Participants viewed rapid streams of pseudo-words with words embedded at regular intervals, while we recorded their EEG. Based on Lochy et al., we expected that words would elicit a steady-state response at the word-presentation frequency (2 Hz) over parieto-occipital electrode sites. However, across 40 datasets (10 participants, two conditions, and two regions of interest - ROIs), only four datasets met the criteria for a unique response to words. This corresponds to a 10% detection rate. We conclude that FPVS should be developed further before it can serve as an individually-sensitive measure of written word processing.

## Introduction

Measures of cognitive processes that are sensitive to individual-level effects can be powerful research tools. Knowledge about individual variance can inform cognitive models and data about individual cognition can have important clinical application. For example, in cognitive neuropsychology, a double dissociation between an individual who can read familiar words but struggles to sound out new words, and an individual with the opposite pattern, gave rise both to the influential Dual Route Cascaded model of reading (Coltheart et al., 2001) and to targeted interventions for distinct reading problems. However, drawing conclusions from individual subjects’ data is often difficult. The influence of measurement error and other sources of “noise” in the data is high in individuals relative to groups, and in clinical applications we often cannot mitigate the noisy single-subject data by testing for long periods.

An emerging method, called ‘fast, periodic, visual stimulation’, or FPVS, is proposed to provide a solution to some of these challenges. It is designed to be a fast way to test cognitive processes in individual subjects with high signal-to-noise ratio (Rossion et al., 2015). In FPVS, participants view rapidly presented stimuli while their neural responses are recorded with electroencephalography (EEG). The stimuli are presented periodically at a certain predictable frequency. Within this stream, we can present stimuli belonging to different categories at different periodic frequencies. For example, we may present scrambled face images every 100 ms (that is, at 10 Hz), with every fifth face presented unscrambled (i.e., unscrambled faces presented every 500ms, at 2Hz). The periodic stimulation is designed to elicit an oscillatory response in the EEG signal at the presentation frequency, in this case at 10 Hz. If the brain also differentiates the embedded or ‘oddball’ category—in this case, the unscrambled faces—that category should also elicit an increased response at the embedded frequency of 2 Hz. Several studies have demonstrated robust responses to the embedded category such as for faces among scrambled face images, for faces among non-face objects, and for new face identities among repeating identities (Rossion, 2014; Rossion et al., 2015). Because the stimulus stream is presented quickly, we can present many stimuli in a few minutes, keeping testing short (approximately 3 minutes). The high signal relative to noise associated with the large number of stimulus presentations is proposed to support individual-level tests of cognitive processes (Liu-Shuang et al., 2014).

The success of these initial face processing studies with FPVS could partly rest on human’s unique face recognition expertise. Recently, researchers have begun to examine whether the same approach can be used to test word recognition. Like face processing, recognising words is a key step in our social and intellectual development. Visual word recognition and face processing are commonly associated with parallel hubs in the left and right fusiform gyri respectively, giving rise to the hypothesis that the visual expertise underlying both processes rests on related mechanisms (McCandliss et al., 2003). In the EEG time course, visual words elicit a larger negative deflection relative to unfamiliar scripts around 170 ms, while face images elicit an analogous ‘N170’ response relative to non-face objects (Bentin et al., 1999). These similarities make word recognition an ideal way to test whether FVPS can be useful in other cognitive domains. On the other hand, evidence that word responses at 170 ms represent any extraction of word meaning is inconsistent, with some studies finding that we can differentiate familiar and unfamiliar scripts, but not the content of words, at that time in the EEG signal (Bentin et al., 1999). Core processing for word meaning appears to emerge around 400 ms from stimulus onset (Kutas & Federmeier, 2011). Further, the point at which we can extract a word’s meaning depends on the word’s length and how frequently we encounter it (Hudson & Bergman, 1985). Thus, stimulus onset for words may be a poorer marker of word processing onset than stimulus onset for faces. Translating FPVS for face processing to word recognition could present some unique challenges.

A study by Lochy et al. (2015) shows promising individual-level effects of word processing using FPVS. They presented words embedded in a stream of scrambled fonts (unfamiliar scripts), nonwords (unpronounceable letter strings), or pseudo-words (pronounceable but meaningless letter strings). Words appeared at 2 Hz in a 10 Hz stream. Words elicited a unique, or “oddball”, response relative to scrambled fonts in 10 out of 10 participants, and relative to nonwords and pseudo-words in 8 out of 10 participants, in under five minutes’ testing for each condition.

These results are exciting because they suggest that FPVS can robustly measure the neural correlates of word recognition on an individual subject level, with minimal testing time. However, individual-level sensitivity could be lower than these headline results suggest. First, fewer people showed an oddball response for words among pseudo-words (8/10), compared to words among scrambled fonts (10/10). This word response among pseudo-words is especially important. Whereas words and scrambled fonts differ in their constituent objects, words and pseudo-words differ in whether the letters form a word. Thus, we can more confidently infer that a person recognises words by looking at their response to words among pseudo-words. Second, this high instance of individual-level word/pseudo-word effects was only reported for the first experiment. In this experiment, neighbouring letters in pseudo-words were not matched to the word stimuli. Particular letter combinations, or ‘bigrams’, are more common than others. For example, in English, ‘st’ is common, while ‘pd’ is rare. Because the first experiment did not match bigram frequency between the words and pseudo-words, the oddball response may have reflected people’s familiarity with letter sequences, as well as, or instead of, recognition of the specific word. The authors tested for a word-specific response among bigram-frequency-matched pseudo-words in a follow-up experiment. They reported a similar group effect but did not report the individual level detection rate, that is, whether the response at the oddball frequency was greater than noise for each participant.

If FVPS can be used to reliably track the neural correlates of word processing in individual subjects, it could have important research and clinical applications. Therefore, we sought to replicate the effect of Lochy et al. (2015), and report the individual-subject detection rate for stimuli matched on bigram frequency. We first established the validity of our implementation by testing whether we could reliably detect faces among scrambled faces. This effect has been replicated many times, so we used it as a sanity check for whether our stimulus presentation and analyses were appropriate. Then, for word processing, we used the design reported in Lochy et al (2015) that produced a high rate of significant effects at the individual-subject level. We used stimulus delivery routines provided by the Rossion group, and the same analysis software and pipeline as that described by Lochy et al. (2015), but used our own stimuli (in English, the original study was in French), participants, and EEG recording system. We asked whether we could replicate their fast, individually-sensitive effects reflecting differential processing of words and pseudo-words. Surprisingly, despite strong individual-subject responses for faces, we found detectable individual-level differentiation of words and pseudo-words in only four out of 40 datasets, suggesting that the method may not be as robust as the initial data suggested.

## Methods

### Participants

We recruited 10 neurotypical adults (seven female, three male) from undergraduate and paid participant pools at Macquarie University. As the study was designed to assess the method at the level of individual subjects, the critical feature for statistical power was the number of trials, rather than the number of participants, so we chose N = 10 to match Lochy et al. (2015). Participants were informed of the aims of the study and gave written consent before taking part. The study was approved by the Macquarie University Human Research Ethics Committee (approval no. 5201200658).

### Stimuli

#### Faces

We used the 50 face images, and 50 scrambled face images from Rossion et al. (2015; available at https://face-categorization-lab.webnode.com/resources/natural-face-stimuli/). The 50 scrambled images had been made by randomly repositioning the pixels in each face image. Consequently, the scrambled faces did not contain coherent object-like features, but had some of the low-level features of faces such as luminance and contrast. It was important for us that the face and non-face stimuli were clearly distinct, as we included this condition to test whether our methods could detect noticeably different stimuli. Because each face image produced a single scrambled face image, scrambled face images repeated four times as often as face images to produce the required ratio. To account for differences in screen resolution between our computer monitor and that used by Lochy et al. (2015), we presented the face and scrambled-face stimuli at twice their original pixel height and width (from 200*200 to 400*400 pixels), and jittered them between 88% and 112% of their mean size. When viewed from 1 m away, the images subtended 6.28 to 8.03 degrees of visual angle in either plane (compared to 5.22 degrees in Rossion et al., 2015).

#### Words (original)

For the word condition, we used 24 common English words and 96 pronounceable pseudo-words (Table S1). Each stimulus was four letters long. We chose to use four-letter words, rather than five-letter words as in Lochy et al. (2015), to maximise the likelihood that the words were acquired early and frequently used. This would make the stimulus set appropriate for fluent readers, and for future use with less fluent readers (e.g., children) or people whose reading ability is unknown. We first chose the 3,000 most imageable words from the Cortese imageability database (Cortese & Fugett, 2004). We entered these words into the Medical College of Wisconsin’s MCWord database to extract word statistics. We sorted the words by most frequently produced, then by imageability. We then used the Oxford Word List, which records word used in Australian children’s writing (http://www.oxfordwordlist.com/, retrieved 15 May 2020), to identify the most frequently written words in Year 2 (typically at age 7-8). This gave us a complementary frequency measure defined by children’s natural word generation. We matched these words with our Cortese word list and selected the 24 items best matched for frequency, imageability, and Year 2 production. We rejected words that were plural (for example, ‘days’) or both a noun and a verb (for example, ‘left’).

We built each pseudo-word by changing the first letter of each target word to another letter that made a pronounceable pseudo-word. We rejected imageable words that could not form appropriate pseudo-words (for example, ‘play’). We replaced these with the next item on our Cortese-Oxford-matched word list. We created four pseudo-words for each word so that each item would appear equally often in our word stream. This should increase experimental control compared to Lochy et al. (2015), as word-specific responses cannot reflect how often they appear in the stream. We submitted each pseudo-word to MCWord to extract word statistics. If a candidate pseudo-word produced a non-zero frequency rating (i.e., was actually a word), we changed its first letter until the frequency rating was zero. We then calculated the bigram frequency mean and standard deviation for each of the five word lists (one word list, four pseudo-word lists). We used a t-test to confirm that there was no evidence for a reliable difference in bigram frequency between each pseudo-word list and the word list (all ps > 0.05, all mean bigram frequencies between 26 and 28). Finally, we used a MATLAB script (MATLAB R2012b, 2012) to generate a JPEG file of each item.

Words and pseudo-words were presented in black Verdana font on a grey background, covering between 1.93 and 6.02 degrees (compared to Lochy et al.’s 3.7 to 6.7 degrees) of the visual field horizontally and between 1.01 and 2.05 degrees (compared to Lochy et al.’s 1.0 to 1.8 degrees) of the visual field vertically when viewed from 1 m with a screen resolution of 1920 x 1080 pixels.

#### Words (large)

Due to different word choices, our words were on average narrower than those in Lochy et al. (2015). To account for the possibility that larger stimuli may be important to elicit clear effects, we also generated large word and pseudo-word stimulus sets by doubling the width and height of our original stimuli. Large word stimuli covered between 3.86 and 12.04 degrees of the visual field horizontally and between 2.02 and 4.1 degrees of the visual field vertically when viewed from 1 m with a screen resolution of 1920 x 1080 pixels.

### Procedure

#### EEG procedure

Data were recorded with a 64-channel BioSemi system sampling at 2048 Hz, using ActiView705 v8.6.1. We first fitted participants with an elastic BioSemi cap and cleaned sites for electromyography with alcohol wipes. We then placed electrodes on the left and right mastoid process, below the right eye, and at the outer canthus of the right eye. Scalp EEG electrodes were attached to the cap in International 10/20 system layout, and connected to the scalp with Signa electrolytic gel. Electrode offsets were below 50 mV.

#### Task

Participants sat 1 m from a computer screen. The task was presented on a 27-inch Samsung S27A950 LED monitor using MATLAB 2012b v8.0.0.783 32-bit (MathWorks, 2012) and Psychtoolbox v3.0.10 (Brainard, 1997; Kleiner et al., 2007; Pelli, 1997). We used presentation scripts provided by the Rossion group which used the same sinusoidal contrast modulation function used by Lochy et al. (2015; SinStim v1.8.9) to fade each stimulus in and out. A fixation cross appeared centrally to mark the start of the trial. The trial began with a random foreperiod between two and five seconds, followed by the stimulus stream (Figure 1). Each stimulus began at 0% contrast, gradually increased to a maximum value, and returned to 0% within 100 ms, creating a stream of stimuli reaching maximum contrast 10 times per second (at 10 Hz). The core of the stream was 60 seconds’ full contrast modulation, in which the maximum contrast was 100% (that is, black stimuli on the white background). The 60 second period was flanked by two-second fade-in and fade-off periods, in which the maximum contrast gradually increased from 0% to 100%, or decreased from 100% to 0%. In each condition, every fifth stimulus was categorically different to the other stimuli; i.e., one face followed four scrambled faces, or one word followed four pseudo-words. This created a 2 Hz stimulation rate for the ‘oddball’ stimuli, faces and words. With 120 unique stimuli presented pseudorandomly to minimise repetitions, this entailed 5 repetitions of each stimulus within the 60 seconds of high contrast stimulation.

**Figure 1.**
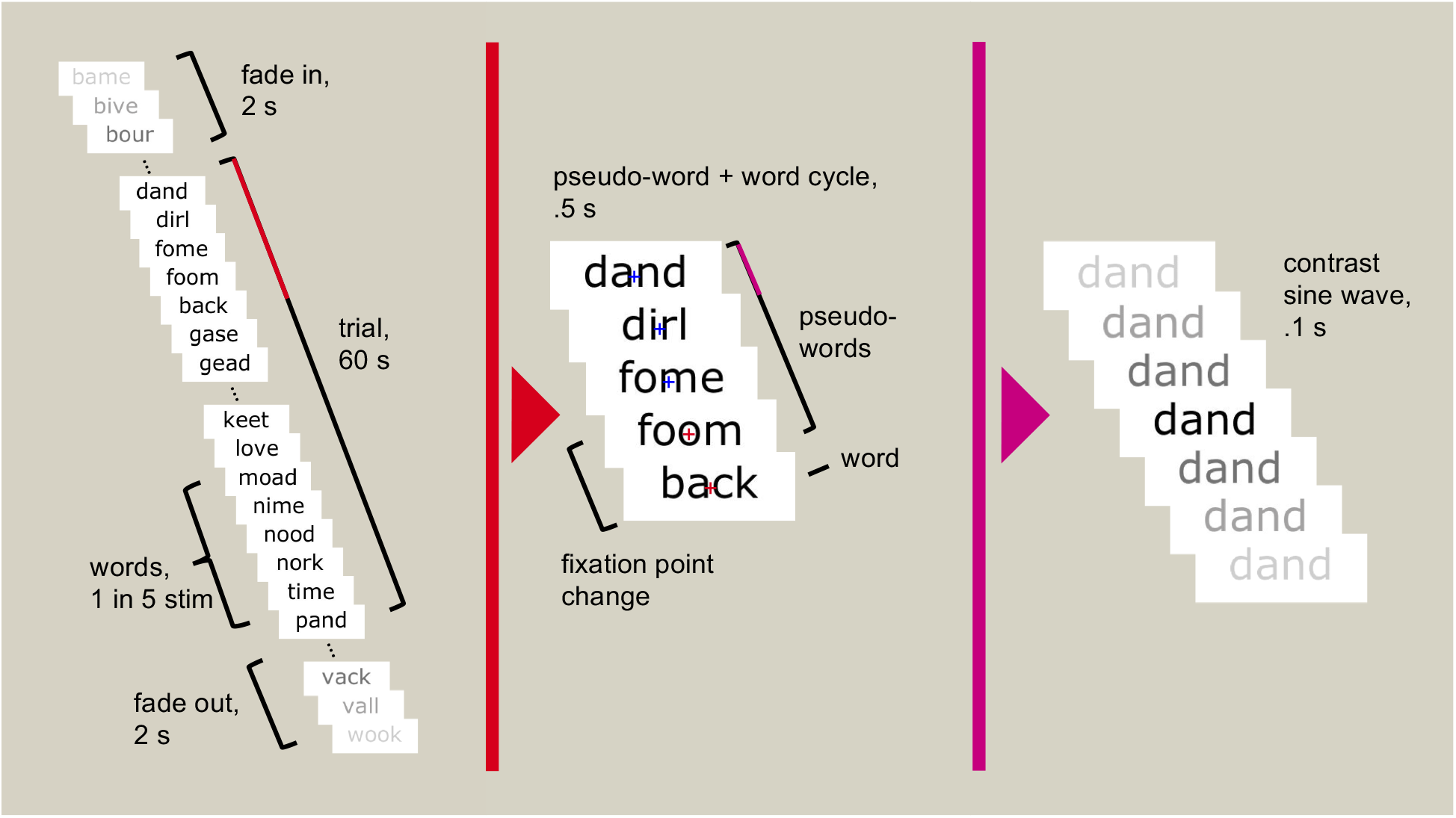
An example trial sequence using words and pseudo-words. Each stimulus reaches maximum contrast and returns to minimum contrast within 100 ms, with a word appearing every fifth stimulus (500 ms, or 2 Hz).

The fixation cross remained centred on screen through the trial. Between six and eight times during the trial, at pseudorandomly-spaced intervals, the fixation cross changed from blue to red for 400 ms. Participants were asked to respond to the change in colour by pressing the space bar on a computer keyboard. The task was designed to ensure that participants maintained focus on the screen without requiring them to engage with the oscillating stimuli.

Participants completed three one-minute trials of each condition: faces and scrambled faces, words and pseudo-words, and large-sized words and large-sized pseudo-words. Condition order was pseudorandomised across participants. We also tested two other conditions for a related project. Testing time was approximately 20 minutes.

## Analyses

### Pre-processing

We followed the analysis steps of Lochy et al. (2015). First, we imported data into Letswave 6 (http://www.nocions.org/letswave/), a MATLAB toolbox used by Rossion et al. (2015) and Lochy et al. (2015) to extract the signal-to-noise ratios of specific frequencies from EEG data. Within this toolbox we processed the Biosemi data with a Fast Fourier Transform filter, width 0.01 to 1 and cut-off 0.1 to 100. We segmented the trials to include fade-in and fade-out periods, creating a 64 s epoch. We then downsampled the data to 250 Hz, linearly interpolated the signal, and re-referenced to the common average. Next, we re-segmented the trials, this time to include only the period of full-contrast stimulation. Because of slight inaccuracies in the monitor refresh rate, stimuli were presented at 10.0007 Hz. Thus, the maximum number of oddball cycles in our 60 s trial was completed at 59.996 s. Following Lochy et al. (2015, 2016), we segmented the trials to this more precise duration. We refer to the stimulation frequency as 10 Hz for simplicity.

Each subject completed three trials in each condition. First, we averaged the data from the 3 segmented trials in each condition together to form a single grand average trial, lasting 59.996s, for each condition. Then, we transformed the grand average data into the frequency domain using a Fast Fourier Transform. We then calculated the signal-to-noise ratio at each frequency of interest by comparing amplitude at that frequency with the 20 surrounding bins (10 bins on each side, each .0166 Hz), excluding the bin with the highest amplitude within these 20 bins. Separately, we calculated z-scores for the frequencies of interest, again relative to the 20 surrounding bins, and excluding one extreme bin. We extracted signal-to-noise ratios and z-scores at both the base (10 Hz) and oddball (2 Hz) stimulation rate for each participant.

### Regions of interest

We expected the base stimulation rate to drive a visual response, so we selected Oz as the electrode of interest. For word and face oddballs (at 2 Hz), we selected symmetrical left- and right-hemisphere regions of interest (ROIs). These were centred on PO7 or PO8 and included the four closest electrode sites (P5, P7, PO3, O1; and P6, P8, PO4, and O2). These scalp locations produced the strongest overall signal-to-noise ratios for the contrast of words and pseudo-words in Lochy et al. (2015). We used their electrode sites *a priori,* rather than focusing on electrode sites with the strongest signal-to-noise ratio (as in Lochy et al (2015)) to avoid possible circularity in the analysis (Kilner, 2013; Kriegeskorte et al., 2009). These sites are also approximately above the fusiform gyrus, which is strongly implicated in both face and visual word processing (e.g., Behrmann & Plaut, 2014). We considered data from left and right hemisphere ROIs separately, as averaging across hemispheres could obscure a lateralised effect.

Faces commonly elicit stronger responses from the right fusiform gyrus, while words typically elicit stronger responses from the left (Maurer et al., 2008; Rossion et al., 2003), which could emerge in our left and right ROIs. However, this lateralisation can vary from person to person, so we did not make specific predictions about where face or word effects should surface for each participant. Following Lochy et al., and in order to be as liberal as possible in replicating the original effect, we did not correct for multiple comparisons across the two ROIs in each oddball analysis. However, assessing two ROIs increases our chance of incorrectly rejecting the null hypothesis (i.e., inflates the false positive rate), so the true effects are likely to be even weaker than we report here (see supplementary material S2 for a deconstruction of our analysis over conditions and ROIs).

### Statistical analyses

#### Group effects

Although our main interest was in individual-subject sensitivity, we first ran a group-level analysis to compare directly with Lochy et al.’s group word-pseudo-word differentiation, and to identify any weak effects that may not appear at the individual level. For each participant, we extracted signal-to-noise ratios at the base rate of 10 Hz, the oddball stimulation rate and its harmonics, 2 Hz, 4 Hz, and 6 Hz. The harmonic at 8 Hz was excluded, following Lochy et al. (2015), who reported that group-level z-scores were not reliably greater than noise at 8 Hz across all conditions. Next, signal-to-noise ratios were averaged across the harmonics, creating two oddball signal-to-noise ratios (left and right ROIs) for each participant in each condition. We collated all participants’ values for each condition and compared them to 1 (the noise level) using Wilcoxon’s signed rank test implemented through the statistical package JASP (Love et al., 2015; compared to the parametric one-tailed t-test in Lochy et al.).

#### Individual effects

For individual-level analysis, we extracted data from Oz at the base stimulation rate (10 Hz), and data averaged across each ROI at the oddball stimulation rate (now the average of 2 Hz, 4 Hz, and 6 Hz). For each frequency of interest we extracted z-scores at that frequency and the surrounding 20 bins. This created two sets of 21 bins per participant per condition: one bin of interest, and 20 bins that in theory should contain no signal, one for the base rate and one for the oddball rate. In Letswave 6, we ranked the bins by their z-score. Following the logic of Lochy et al., if there was no effect (similar response in all bins), each bin would have an equal 5% chance of ranking first. Thus, allowing for a 5% false positive rate, we concluded that any participants whose strongest z-score was at the oddball frequency showed an oddball-specific response.

## Results

### Behaviour

All participants responded to fixation point changes with a hit rate of 70% or higher on each trial.

### Visual steady state response (10 Hz) at Oz

We first checked for responses driven by visual stimulation at the base frequency of 10Hz, in the occipito-central electrode (Oz).

#### Group analysis

The base stimulation frequency was reliably greater than noise in the face condition and both word conditions (Table 1).

**Table 1.** Descriptive and inferential statistics for visual steady state responses (SNR) at 10Hz, reflecting the base stimulation frequency (visual stimulation). Wilcoxon’s V indicates the sum of the positive differences between observed and test values. Large values, relative to the number of observations, indicate a larger difference.

|  | Median | SD | Wilcoxon's V | p |
| --- | --- | --- | --- | --- |
| Faces in scrambled faces | 22.10 | 18.03 | 55.00 | < .01 |
| Words in pseudo-words, original | 8.01 | 12.70 | 55.00 | < .01 |
| Words in pseudo-words, large | 8.57 | 10.82 | 55.00 | < .01 |

*Note.* Degrees of freedom = 9.
*Note.* Hypothesis: median signal-to-noise ratio is greater than the noise level, 1.

#### Individual-level analysis

The base stimulation frequency also produced detectable individual-level effects (p < 0.05), as defined by Lochy et al. (2015) and described above, at Oz. This was the case for every participant in every condition, with two exceptions. One participant did not meet the criteria for a detectable effect in the word condition with large stimuli, but did in the original-sized word condition. Another participant showed the opposite pattern. All participants met our criteria for a visual steady-state response in at least one condition. We include individual-level frequency spectra for each condition in Supplementary Material S3.

### Oddball (category) response for faces (2 Hz)

Next, we checked whether we could detect 2Hz oscillatory responses reflecting the presentation of faces among scrambled faces, in our left and right ROIs centred on P07 and P08.

#### Group analysis

As expected, oddball faces elicited a statistically reliable effect at 2 Hz in both the left and right ROI, as measured by Wilcoxon’s signed rank test (Table 2; see Figure 2 for frequency spectra).

**Figure 2.**
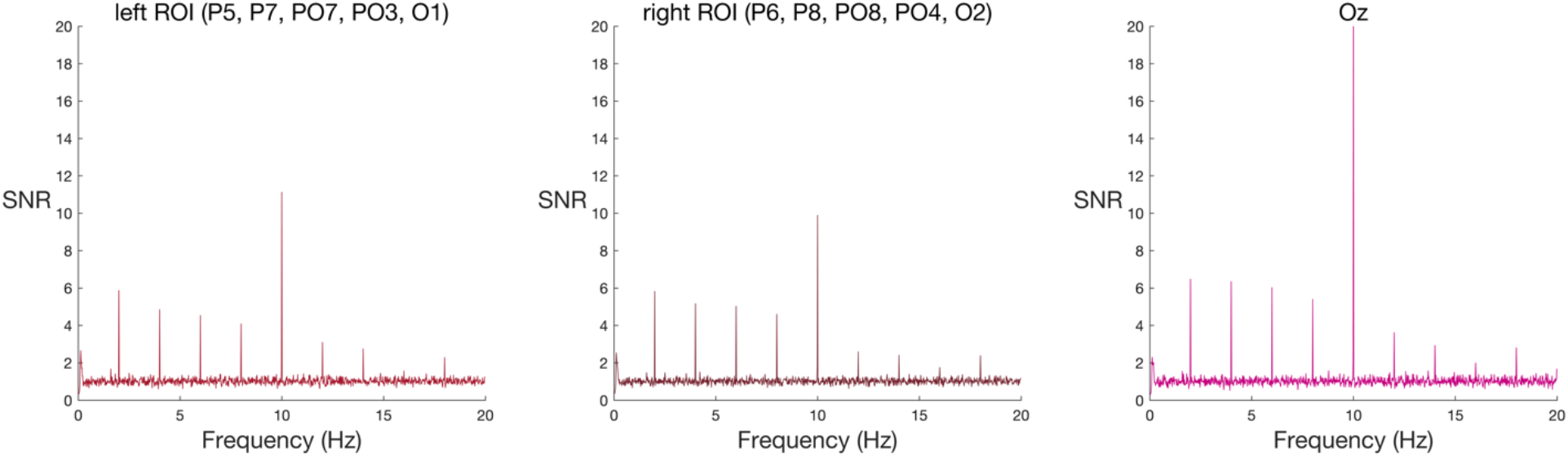
Signal-to-noise ratio frequency spectra across the group for faces among scrambled faces. Visual steady state responses appear at 10 Hz. Face-specific responses appear at 2Hz and its harmonics (4Hz, 6Hz, 8Hz).

**Table 2.**
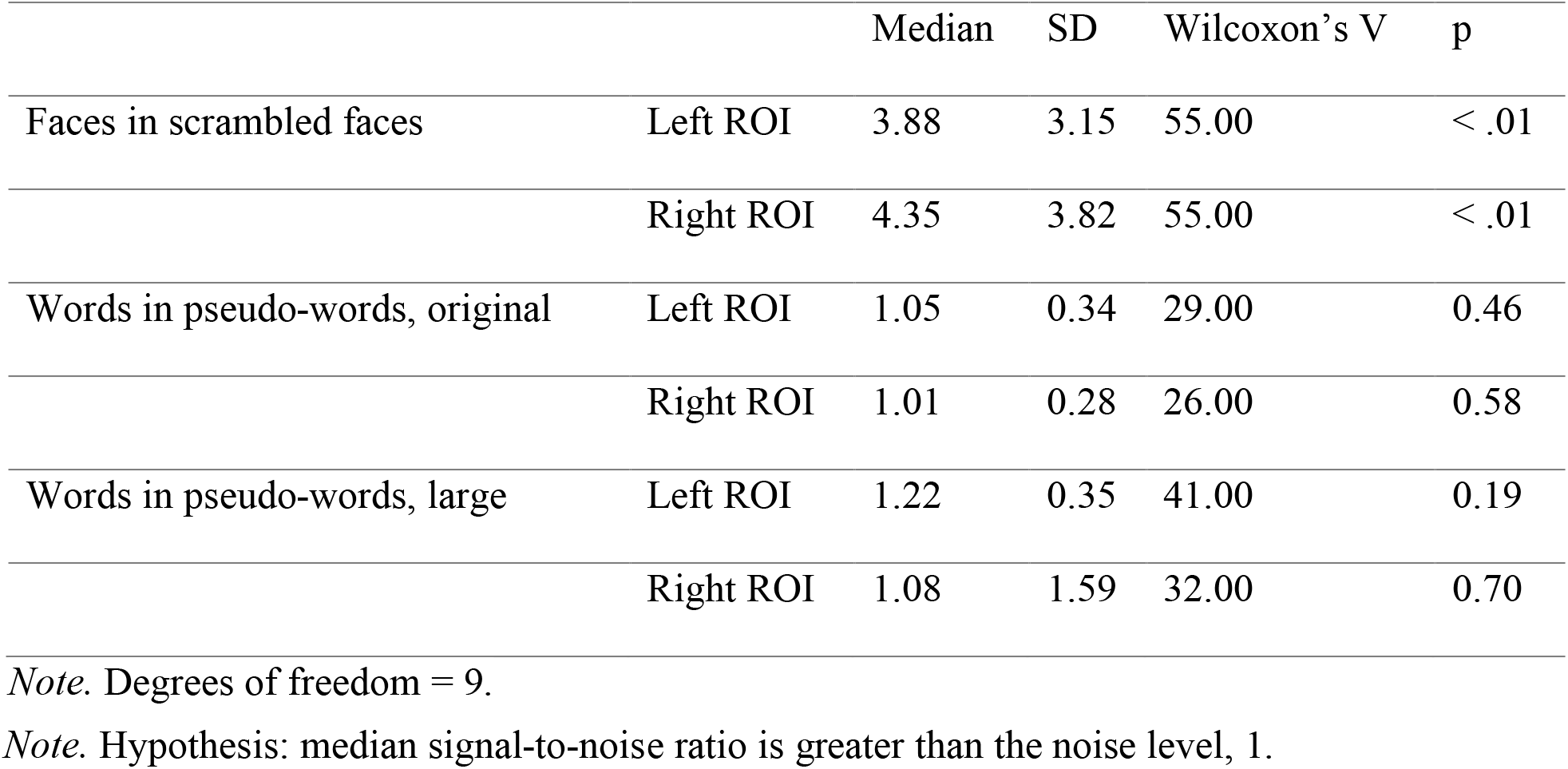
Descriptive and inferential statistics for stimulus differentiation (SNR) at 2Hz.

#### Individual-level analysis

In line with findings across the group, most (9/10) participants met the criteria for a face-specific response. These nine participants showed the effect in both ROIs. The remaining participant’s strongest signal-to-noise ratio in the sampled range did not fall on the face frequency in either the left or the right ROI.

### Oddball (category) response to words (2 Hz)

Finally, we asked whether we could replicate the main result of interest: a 2Hz oscillatory response reflecting the presentation of words among pseudowords.

#### Group analysis

In contrast to the robust face effect, and counter to the outcome of Lochy et al. (2015), neither set of words reliably elicited a response greater than noise at the group level (Table 2; see Figure 3 for frequency spectra).

**Figure 3.**
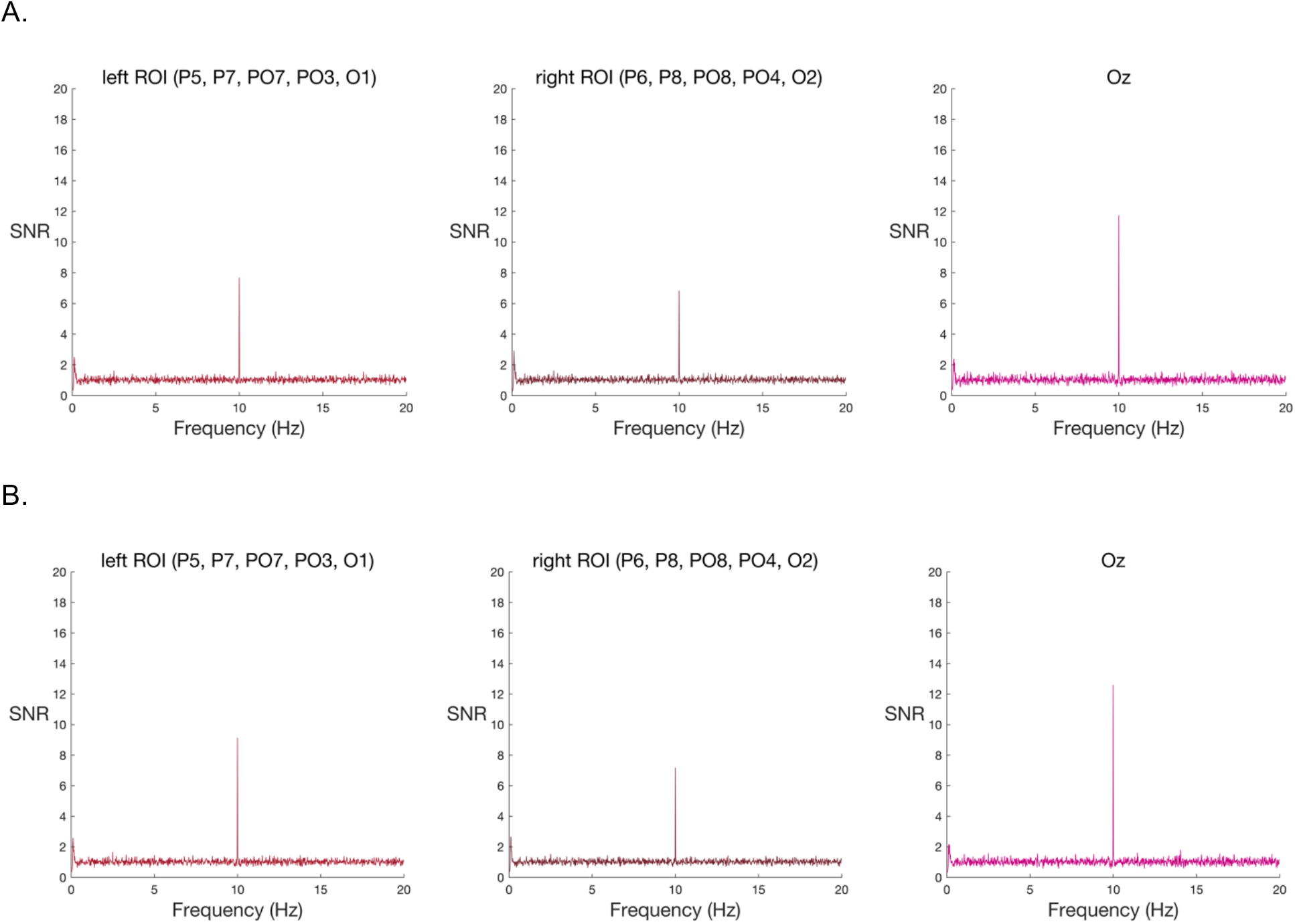
Signal-to-noise ratio frequency spectra across the group for (A) original-sized and (B) large words among pseudo-words. Visual steady state responses appear at 10 Hz. Word-specific responses were analysed at 2Hz and its harmonics (4Hz, 6Hz, 8Hz).

#### Individual-level analysis

Again, in contrast to the face effect and the results of Lochy et al. (2015), but consistent with our failure to find significant effects at the group-level, individual word-specific responses were rare. For original-size stimuli, only one participant showed a word-specific response and only in the left ROI. For large stimuli, two participants showed a word-specific response; one in both ROIs, and the other only in the left ROI. In total, across four tests—two word-stimulus sizes and two ROIs— and 10 participants (a total of 40 datasets) only four datasets met the criteria for a unique response to words nested among pseudo-words, using an uncorrected alpha level of 5%. This corresponds to a 10% detection rate (see Figure S3). When we corrected the alpha level, to control for the 4 multiple comparisons inherent in testing 2 ROIs and 2 stimulus sizes in each person, none of the datasets showed a significant effect (see Table S2).

## Discussion

We set out to replicate the claim that fast, periodic stimulation could provide an individually-sensitive, minimally-demanding test of word recognition. Previous work suggests that FPVS can rapidly elicit a unique response to words embedded among pseudo-words in 8 out of 10 people (Lochy et al., 2015). We asked whether word detection rates would be similarly high in a novel sample of English speaking adults with word and pseudo-word lists matched for bigram frequency. We expected to observe similarly high word-detection rates in adults who we knew were able to read, with the eventual aim of using the test for clinical purposes in the future.

Contrary to our prediction and the previous literature, we observed very low detection rates for neural entrainment to words embedded in pseudo-words. After validating our stimulus delivery and analysis pipeline using face stimuli (near perfect detection rate – 18/20 datasets) we tested word detection in two ROIs (as in Lochy et al.) with two stimulus sizes. Of the 40 word datasets, only four showed a statistically reliable effect, corresponding to just a 10% detection rate, even when using an uncorrected alpha of 5%.

Previous studies highlight reasons that word processing can be more difficult than face processing to track in the EEG trace. Particularly important for our study is the finding that rapid responses to words (around 170 ms from onset) can be driven by the visual characteristics of words, rather than their meaning (Maurer et al., 2008). Even in Lochy et al. (2015), we see that word detection drops when words are embedded in pseudo-words formed from real letters, instead of false fonts.

These rates could have dropped further in our study as our words and pseudo-words shared more statistical features. However, the only essential difference between the studies was that we matched our words and pseudo-words for bigram frequency. Considering how much lower detection rates are in our sample, it is surprising that bigram frequency should make such a difference. Bigram frequency is typically controlled in studies of early word processing, leaving it unclear how strong an effect it might have. In line with standard practice, Lochy et al. control for bigram frequency in a secondary study in their 2015 paper. They do not report individual effects for this analysis, but report no major change in the group effect, whereas we could not detect an effect in the group effect.

Another explanation for the disparity in detection rates could be that the limited sample size in both studies did not fully capture the distribution of responses. We limited our sample size to that of Lochy et al., anticipating that most participants would show a word-specific response. This drop in single-subject detection rates is sufficient to conclude that word detection with FPVS is not always as robust as one might anticipate given Lochy et al.’s 80% success rate, but leaves us unsure how much random variation exists across individuals in their neural responses to words. If word FPVS only elicits word-specific responses in some people, this would fundamentally limit how we use the method to study word processing in single subjects. Clarifying how the design (e.g., whether words are embedded among pseudo-words with the same or different statistical features) and individual variation in brain responses drive detection rates will be important for understanding how FPVS can best be used.

In conclusion, fast, periodic stimulation promises rapid insight into mental processes with strong signal relative to noise. Despite many examples of individual-level face detection with FPVS, and preliminary evidence that FPVS can elicit word-specific responses in single subjects, here we showed that individual-level word detection is difficult to probe even with FPVS. We suggest that FPVS needs to be developed further before it can serve as a single-subject measure of written word processing.

## Supplementary Materials

S1. Word and pseudo-word lists

**Table S1.**
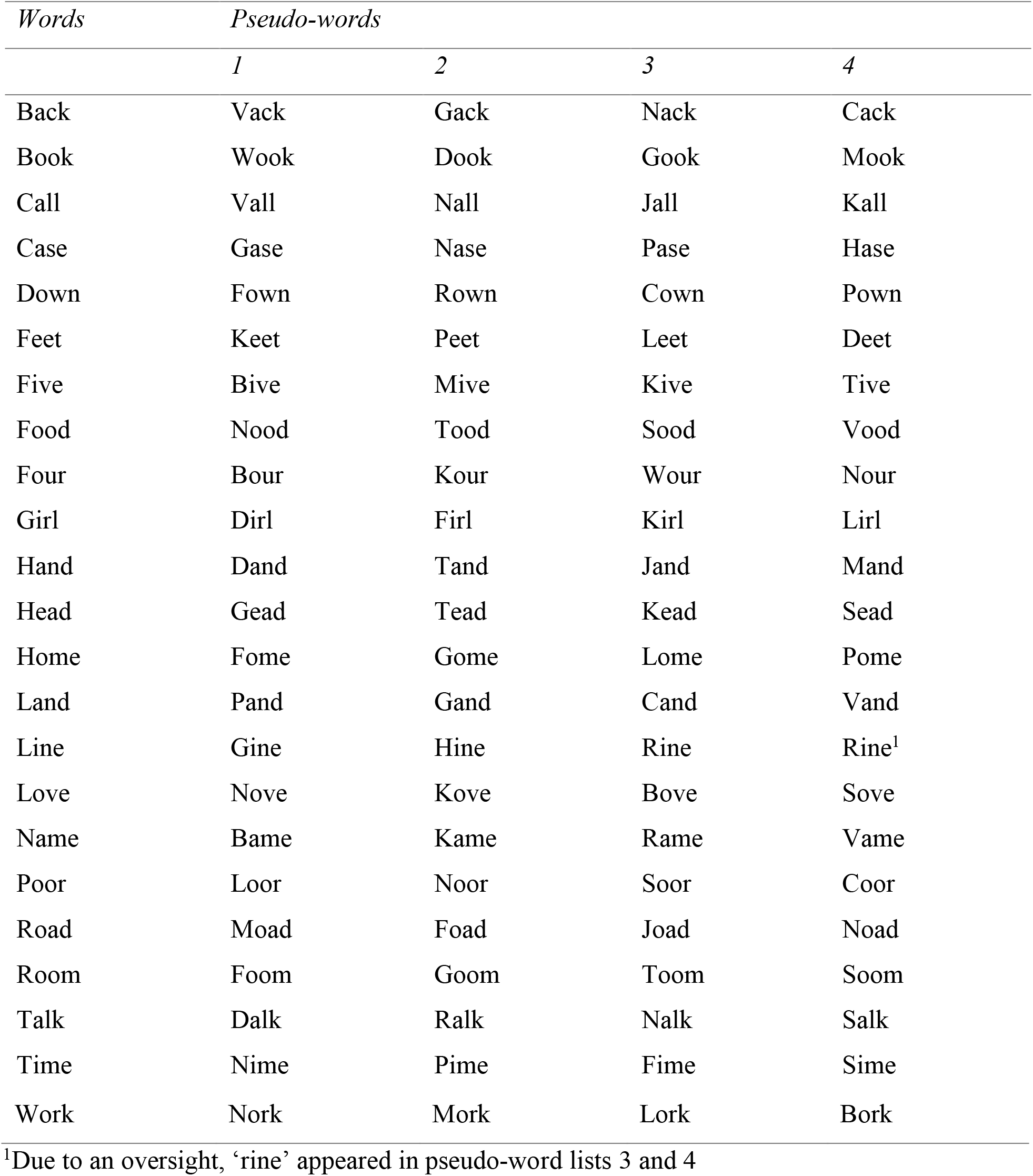
The first letter of each word was changed four times to create four pseudo-words. Each pseudo-word list was then compared to the word list for bigram frequency.

S2. Word-specific response detection rates adjusted for multiple ROIs and stimulus sets.

We tested whether people showed a word-specific response in four different ways: with two stimulus sizes, in two regions of interest. We did this to sensitively test whether we can elicit word-specific responses in a high proportion of people. However, for transparency, here we show the detection rates after correction for the number of tests we conducted.

First, we show the number of individuals who met the standard criteria for a word-specific response. We break it down by condition to show how many would have met those criteria had we limited our study to one ROI and one stimulus set. For the most successful combination, if we were to use only large stimuli and exclusively looked at the left ROI (the typical site of language lateralisation), our detection rate would be 2 out of 10 people (20%).

Second, we recompute the z-scores we used to judge detection rates. The original method asks whether the frequency of interest (2 Hz and its harmonics) is larger than the 20 surrounding frequency bins. This should only happen by chance 5% of the time. We reran this analysis, selecting the surrounding 40 and 80 bins to establish 2.5% and 1.25% false positive rates. This corrects for two (stimulus sets) or four (ROIs*stimulus sets) comparisons. The frequency resolution was .0166 Hz, meaning that 80 bins around 2 Hz, excluding the first neighbouring bins, captured the range from 1.30 Hz to 2.70 Hz. When we control the false positive rate at 1.25%, no participant meets the criteria for a word-specific response in any condition.

**Table S2.** Number of individuals (out of ten) who met the criteria for a word specific response in each condition, controlling the false positive rate at 5%, 2.5%, and 1.25%

|  | *Small stimuli* | *Large stimuli* |
| --- | --- | --- |
| *alpha = .05* |  |  |
| *Left ROI* | 1 | 2 |
| *Right ROI* | 0 | 1 |
| *alpha = .025* |  |  |
| *Left ROI* | 1 | 2 |
| *Right ROI* | 0 | 1 |
| *alpha = .0125* |  |  |
| *Left ROI* | 0 | 0 |
| *Right ROI* | 0 | 0 |

**Figure S3.**
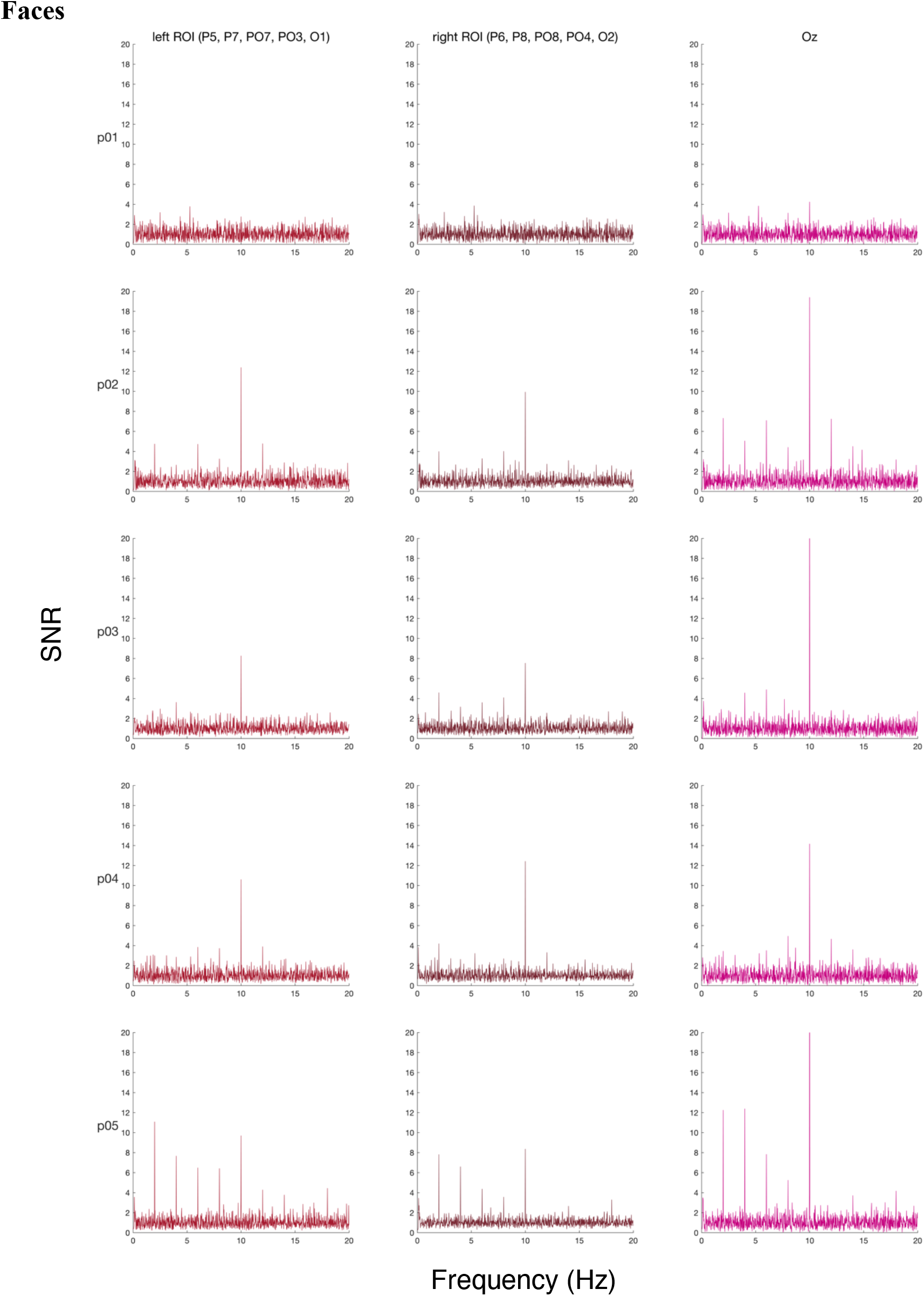
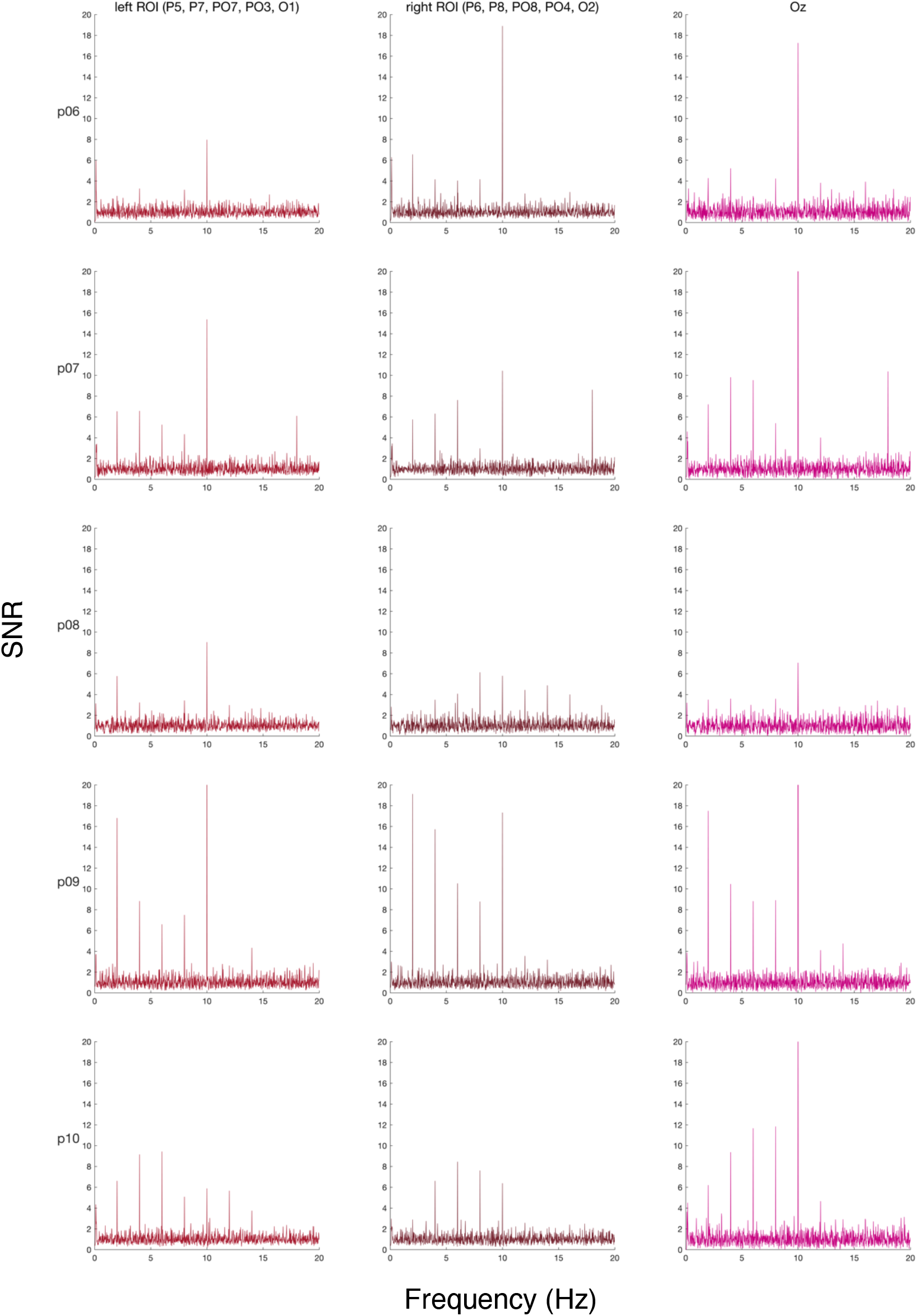
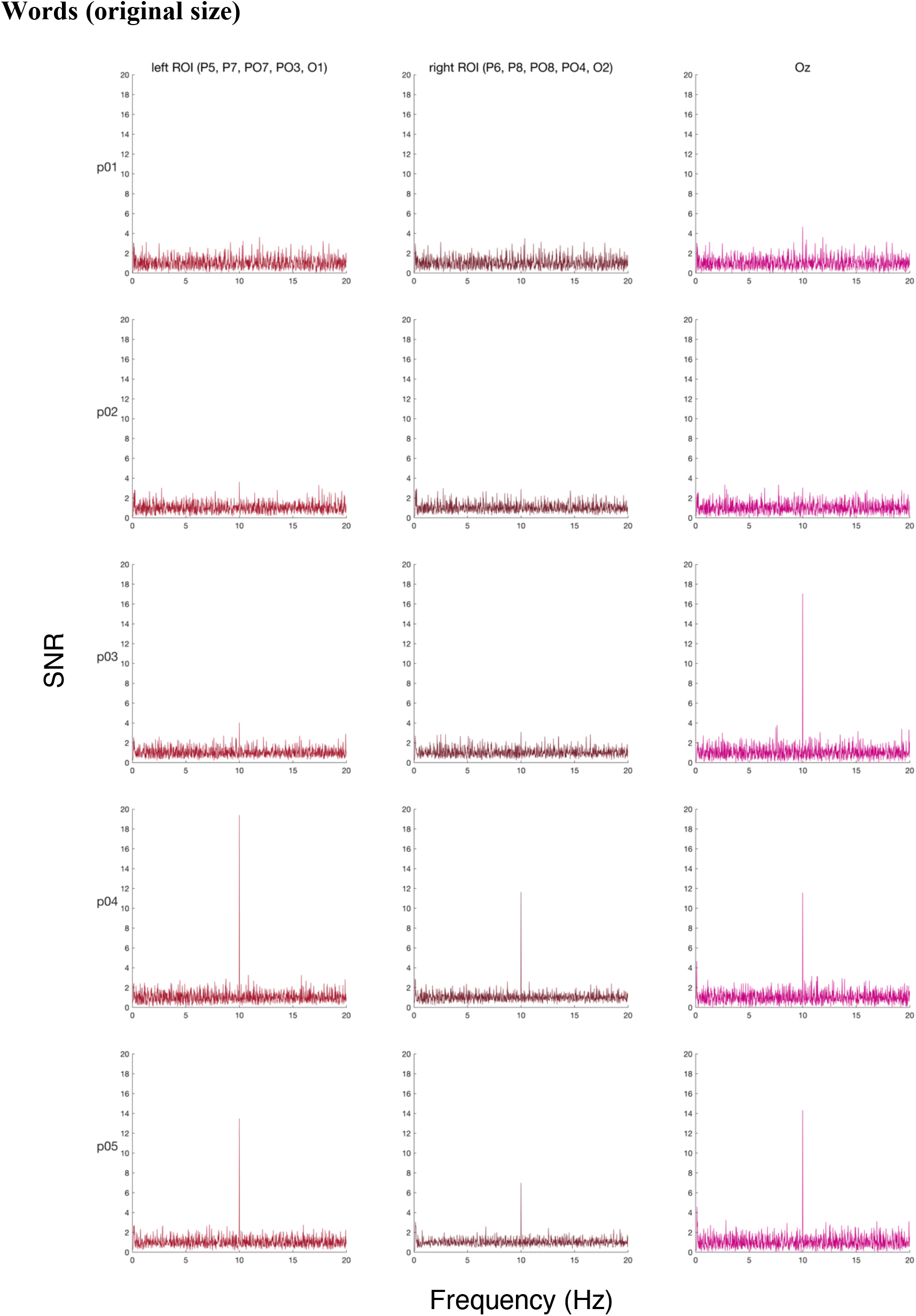
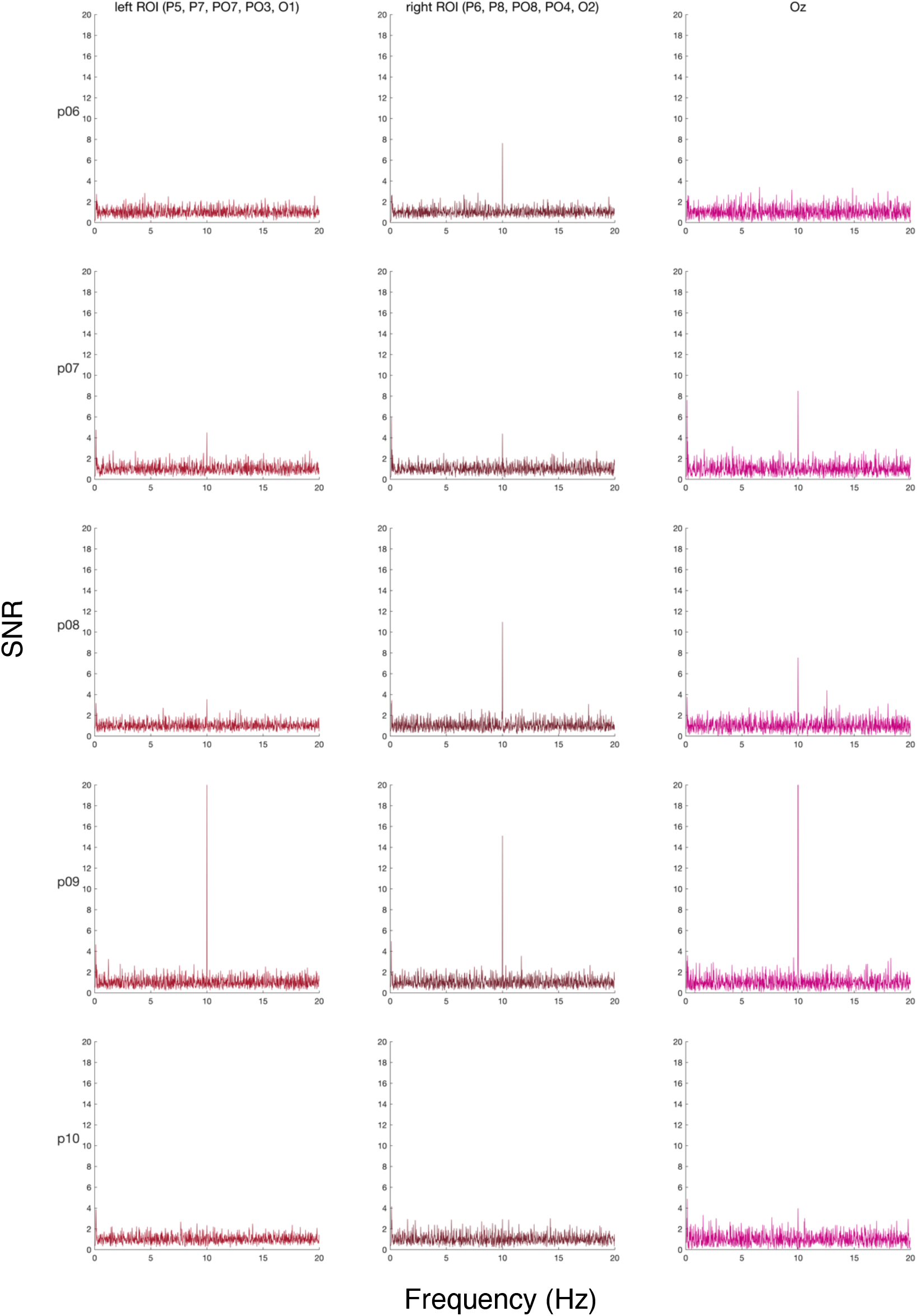
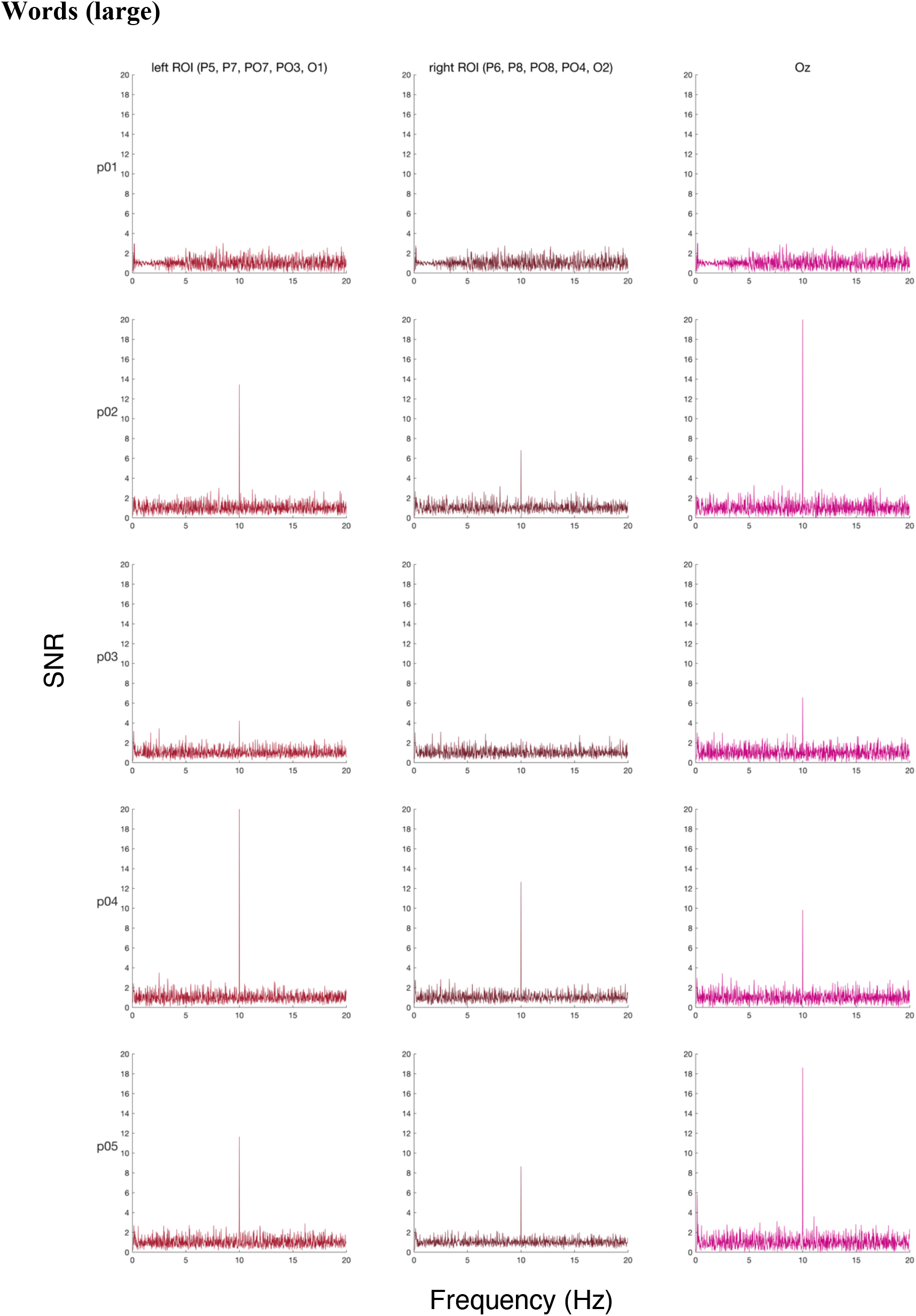
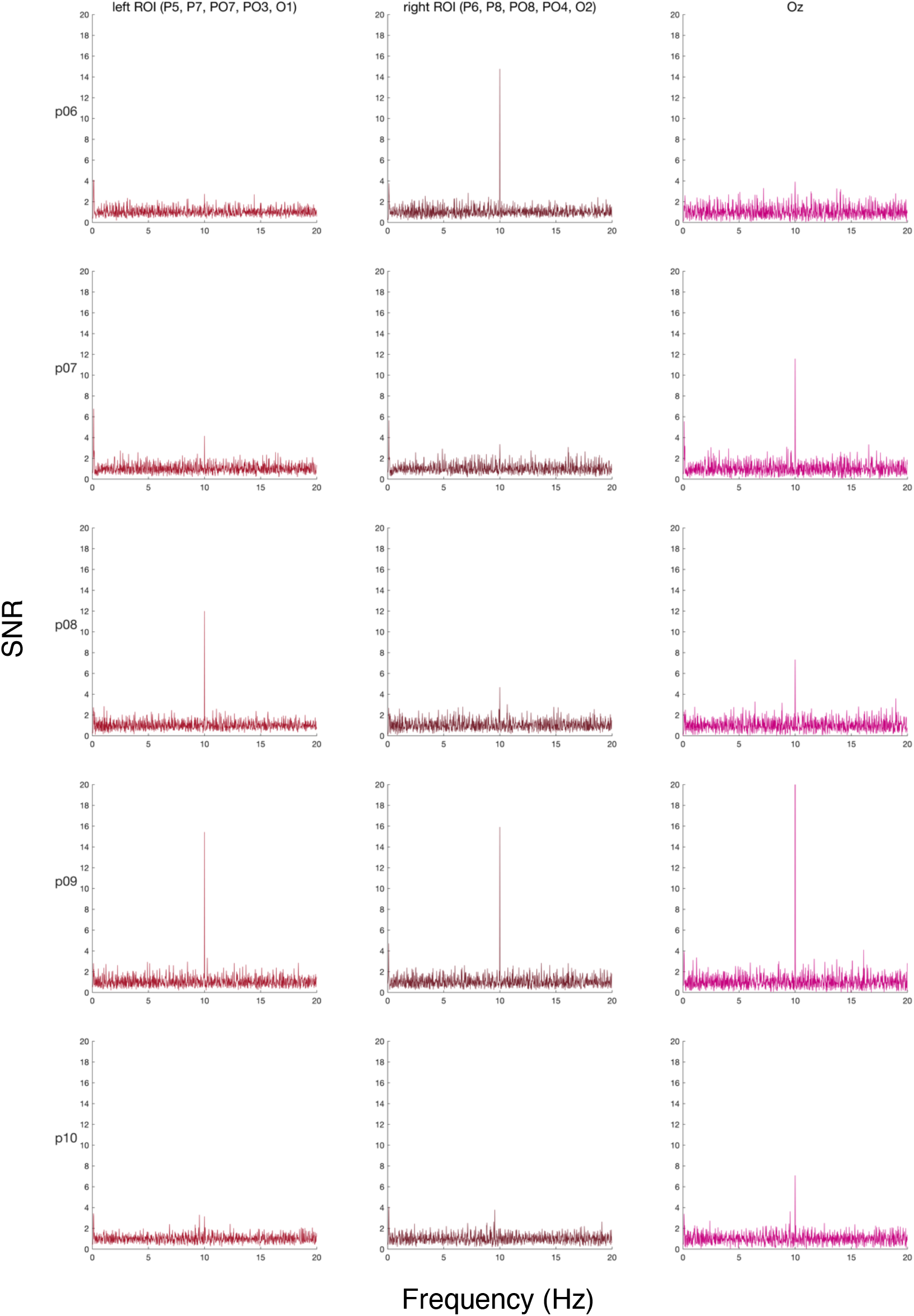
Individual participant signal-to-noise ratio frequency spectra for face, word (original), and word (large) conditions over left, right, and central ROIs.

